# Ab initio phasing macromolecular structures using electron-counted MicroED data

**DOI:** 10.1101/2021.10.16.464672

**Authors:** Michael W. Martynowycz, Max T.B. Clabbers, Johan Hattne, Tamir Gonen

## Abstract

Structures of two globular proteins were determined *ab initio* using microcrystal electron diffraction (MicroED) data that was collected on a direct electron detector in counting mode. Microcrystals were identified using a scanning electron microscope (SEM) and thinned with a focused ion-beam (FIB) to produce crystalline lamellae of ideal thickness. Continuous rotation data were collected using an ultra-low exposure rate on a Falcon 4 direct electron detector in electron-counting mode. For the first sample, triclinic lysozyme extending to 0.87 Å resolution, an ideal helical fragment of only three alanine residues provided initial phases. These phases were improved using density modification, allowing the entire atomic structure to be built automatically. A similar approach was successful on a second macromolecular sample, proteinase K, which is much larger and diffracted to a modest 1.5 Å resolution. These results demonstrate that macromolecules can be determined to sub-Ångström resolution by MicroED and that *ab initio* phasing can be successfully applied to counting data collected on a direct electron detector.

## Main

The cryogenic electron microscopy (cryoEM) method microcrystal electron diffraction (MicroED) can be used to determine the atomic structures of inorganic materials, small organic molecules, natural products, soluble proteins, and membrane proteins from vanishingly small crystals^1^. The data are collected as a movie on a fast camera while the stage of the electron microscope is continuously rotating the crystal in a parallel electron beam^2^. Since the method is analogous to the rotation method used in X-ray crystallography^3^, data processing is conducted using standard X-ray crystallography software. For MicroED of small molecules and short peptides which typically diffract to atomic resolution, phases are commonly determined using *ab initio* direct methods^4^. Phases for electron diffraction can also be determined from images recorded on a transmission electron microscope (TEM)^5–7^, as demonstrated for two-dimensional membrane protein crystals. For MicroED data from three-dimensional macromolecular crystals, phases have thus far only been determined by molecular replacement^1^. Molecular replacement for MicroED data has been demonstrated with distant homologues^8,9^ or extracted fragments from homologues^10^, but there have been no reports of successful phasing in the absence of a previously known model. A major obstacle for phasing using idealized fragments in macromolecular MicroED data has been phase improvement^10^; to date, density modification algorithms have persistently failed to improve the maps for any MicroED data from macromolecules. Thus, even if idealized fragments had been placed accurately, phasing was intractable. Robust phasing by any means other than molecular replacement has remained elusive for macromolecular MicroED^1^.

MicroED data have been collected using a variety of camera systems and microscopes. While a charge-coupled device (CCD) can be used^11^, these cameras tend to be slow and relatively insensitive, which makes data collection by continuous rotation difficult. Cameras based on complementary metal-oxide-semiconductor (CMOS) technology allow faster readout and often have better signal-to-noise ratios^1,12^. The shorter dead time between frames allows CMOS cameras to operate in rolling-shutter mode, which facilitates continuous rotation experiments^2^. Hybrid pixel detectors (HPDs) have also been successful for collecting macromolecular MicroED data^9,13.^ HPDs have a smaller array of large pixels that can complicate data collection when working with large unit cells such as proteins. In contrast, direct electron detectors offer distinct advantages and were pivotal for the “resolution revolution” in single-particle cryoEM^14,15^. In electron-counting mode, these low-noise cameras can resolve individual electron events. The increased accuracy of electron counting in combination with faster readout rates promises to deliver superior data quality for MicroED. Direct electron detectors have been used in prior MicroED investigations, but only in integrating (linear) mode^16,17^. It was believed that the number of electrons in the diffraction spots was too high to reasonably count given the low frame rate of the detector^16^.

Here, a structure of triclinic lysozyme at 0.87 Å using a Falcon 4 direct electron detector in electron counting mode is reported (**Figure 1**). The exposure rate, frame rate, and total exposure time of the MicroED experiment were optimized to allow data collection within the specifications of the camera (**Materials and Methods, Supplementary Figure 1**). Using this approach, far superior data was collected when compared with previous studies, and phases were determined *ab initio* using an idealized fragment of only 15 atoms. Initial phases from these atoms were improved by density modification^18^, resulting in a map showing individually resolved atoms for essentially the entire protein. A model was built automatically into this map using standard crystallographic software^19^ without user intervention. To test the robustness of this approach, data were collected from crystals of the serine protease proteinase K (**Supplementary Figure 2**). The data extended to 1.5 Å resolution with better statistics than previously reported structures of this protein using other detectors^20,21^. This structure was initially phased by automatically placing four ideal helices followed by successive rounds of density modification and backbone tracing. The backbone model was then completed by automated means. This study demonstrates electron-counting MicroED data collection to sub-Ångström resolution using a direct electron detector. Even at near-atomic resolutions, density modification could be successfully applied, leading to *ab initio* phasing of macromolecular MicroED data. These structures set a new benchmark for achievable quality of MicroED data and increase the scope of possible experiments targeting proteins with unknown structures.

**Figure 1.**
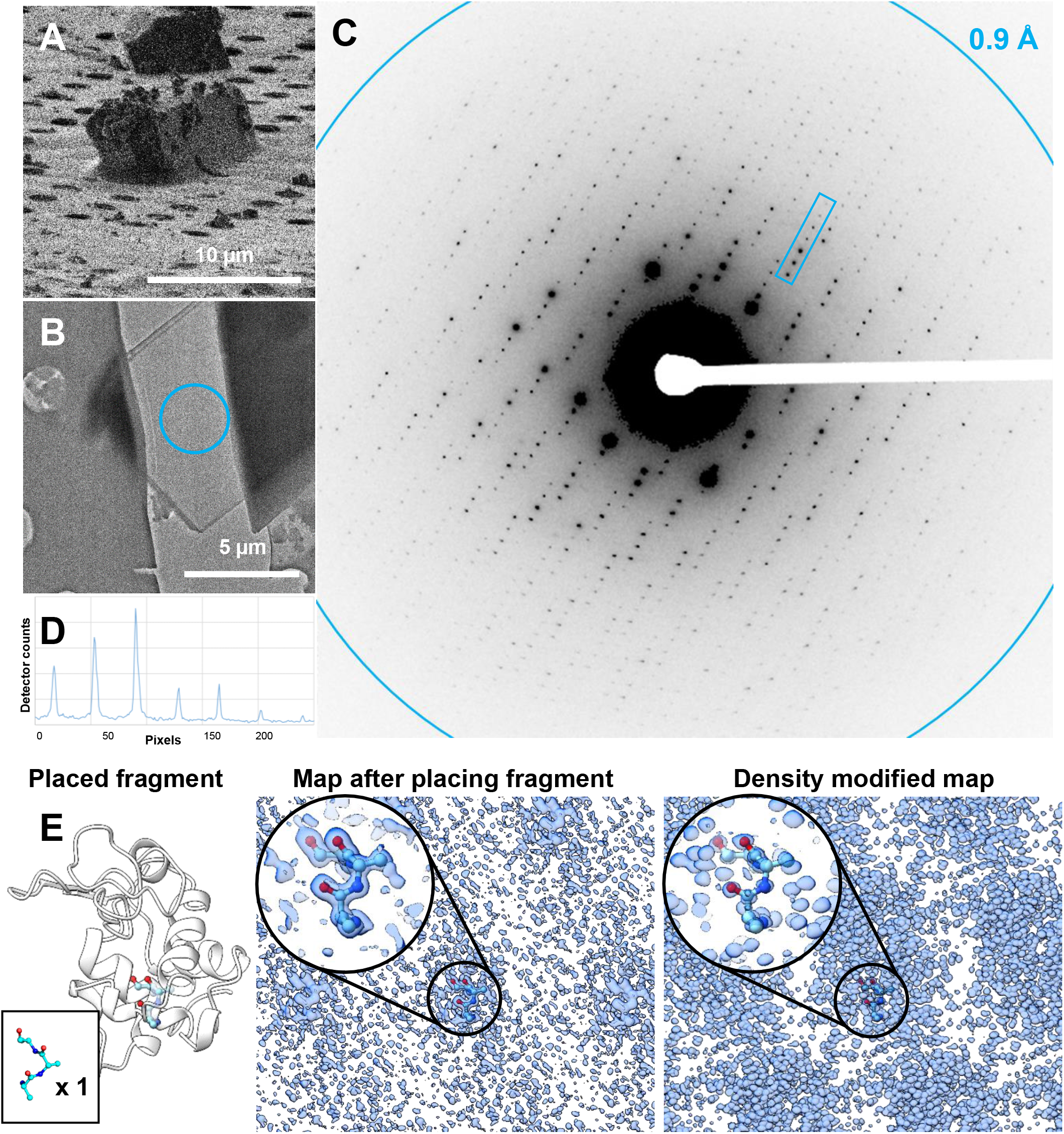
Electron counted MicroED data from milled lamellae of lysozyme. (**A**) A typical lysozyme microcrystal imaged using the focused ion beam. **(B)** A thin, milled lamella from (A) identified in the TEM. **(C)** MicroED data collected in counting mode using a direct electron detector extending to atomic resolution. The data were summed over a two-degree wedge for display purposes only. **(D)** A line plot through the box indicated in (C) is inset demonstrating the quality of the counting mode data collected showing high I/σ. (**E**) The ab initio phasing strategy where a small fragment was placed, and the initial phases were extended by desnity modification. The position of the placed fragment and the maps before and after density modification are displayed as indicated to the right.

Crystals of triclinic lysozyme were grown in batch (**Materials and Methods**). The crystals were visible under a light microscope and were initially about 10 μm in size. A slurry of these microcrystals were vitrified on EM grids. Grids were loaded into a dual-beam FIB/SEM where individual crystals that were at least 5 μm from a grid bar and at least 3 grid squares away from the edge of the grid were identified. The crystals were then coated in multiple layers of platinum for protection from the ion beam during the milling process. Each crystal was then milled into a thin lamella approximately 300 nm thick, or about the ideal thickness for MicroED data collection at 300 kV accelerating voltage based on prior mean free path measurements^22^ (**Figure 1A, B, Materials & Methods**).

The milled lamellae were then loaded under cryogenic conditions into a Titan Krios. Lamellae sites were identified by low-magnification montages, where each site appeared as a semi-transparent shape suspended above a strip of empty area. 18 out of the 20 milled lamellae were found in the TEM-two did not survive the cryotransfer step. For each lamella, the eucentric height was carefully adjusted. Each lamella was inspected initially at an intermediate magnification to verify that no contamination had built up or compromised the sample (**Figure 1B**). Each lamella was checked for diffraction and the camera length was adjusted appropriately depending on the attainable resolution before data collection.

MicroED data was collected in electron counting mode using a Falcon 4 direct electron detector (**Materials and Methods, Supplementary Figure 1, Figure 1C, D**). In counting mode, this camera allows for highly accurate detection of single electron events. However, the number of electrons that can accurately be counted in each pixel of each frame is limited. To assure accurate reporting of the intensities, the exposure rate must be kept very low. This strategy reduces errors caused by too many electrons hitting the same pixel within a readout cycle of the detector, but risks missing weak reflections in the background. These stringent camera requirements could be met by greatly reducing the exposure rate and compensating by increasing the total exposure time (**Materials and Methods**). This strategy prevents the strong reflections from overwhelming or damaging the detector while sampling the weak reflections at high resolution was achieved at sufficient frequency to recover their intensities (**Supplementary Figure 1**).

Multiple settings had to be adjusted to achieve a suitably low exposure rate for these experiments in electron counting mode. Importantly the camera’s dose protector, which automatically retracts the camera when the microscope enters diffraction mode, must be disabled (**Materials and Methods**). The smallest C2 aperture was used coupled with the highest spot size possible on our instrument, 50 μm and spot size 11, respectively. The instrument was kept in microprobe mode to avoid an approximately fivefold increase in exposure rate that occurs by deactivating the condenser mini lens. Since these experiments were conducted on a Titan Krios, the beam size could be changed while maintaining a near-perfect parallel illumination. The beam diameter was spread to 25 μm to reduce the exposure per unit area further. Together, these modifications reduced the total exposure per area by a factor of up to 10 × compared to prior experiments on this instrument^23^. Further, the data was collected over a 420 s exposure time at 250 frames per second, which was the longest possible on the detector software, coupled to a very slow rotation rate of the stage at either 0.15 or 0.2 °/s, corresponding to a total real space-wedge of approximately 60° or 80°, respectively (**Figure 1C**). In this way the exposure was spread in space and time, allowing accurate measurements of single electron events even in diffraction mode. As a result, these datasets had total exposures up to 4 × lower than prior investigations for similar macromolecules even though the recording time here was more than twice as long^24^. The total exposure per lamella was ∼0.64 e^-^/ Å^2^. The background noise and total flux on the camera were further minimized by using a 100 μm selected area aperture that corresponds to a region about 2 μm in diameter on the specimen. Under these conditions, essentially all pixel values fall within the linear range of the detector and all are well below the threshold for damaging the detector (**Materials and Methods**).

The movies from the Falcon 4 were sliced into 0.5 s segments, or 840 individual frames, each 2048×2048 16-bit pixels in size, spanning 0.075°–0.08° rotation. Images were converted to SMV format using the MicroED tools^12^ adapted for the Falcon 4/EPU-D metadata format. The total size of a compressed MicroED movie in counting mode for these exposures is typically 2.2 GiB, regardless of the number of frames (up to 14,455 in 420 s). The size of these movies reflects the total information content captured. The converted frames were processed using standard crystallographic software^25,26^. The space group for all 18 lysozyme MicroED datasets were found to be *P* 1, and the typical unit cell was determined to be (a, b, c,) (Å) = (26.42, 30.72, 33.01), (α, β, γ) (°) = (88.319, 109.095, 112.075) (**Table 1**). The data were merged to increase completeness. The subsequent merging steps identified two lysozyme lamellae that correlated poorly with the other 16 integrated datasets. These two datasets were discarded. A high-resolution cutoff at 0.87 Å was applied. This corresponds to where the CC_1/2_, the correlation coefficient between randomly chosen half-datasets^28^, was significant at the 0.1% level. The overall completeness of the final dataset was 87.5%, with all resolution shells below 1 Å resolution being > 95% complete (**Table 1, Supplementary Table 1, and Supplementary Figure 3**).

**Table 1.** MicroED crystallographic table of triclinic lysozyme.

| MicroED Data and Structure |  |
| --- | --- |
| Triclinic Lysozyme |  |
| Wavelength (Å) | 0.0197 |
| # of crystals (#) | 16 |
| Resolution range (Å) | 16.05 - 0.87 (0.9011 - 0.87) |
| Space group | P 1 |
| Unit cell (a, b, c) (Å) | 26.42, 30.72, 33.01 |
| ($\alpha$, $\beta$, $\gamma$) (°) | 88.319, 109.095, 112.075 |
| Total reflections (#) | 569407 (5797) |
| Unique reflections (#) | 64986 (2783) |
| Multiplicity | 8.8 (2.1) |
| Completeness (%) | 87.55 (37.64) |
| < I/$\sigma$(I) > | 6.23 (0.66) |
| Wilson B-factor | 9.44 |
| R-merge | 0.2363 (1.035) |
| R-meas | 0.248 (1.409) |
| R-pim | 0.0730 (0.9451) |
| CC<sub>1/2</sub> | 0.99 (0.147) |
| CC* | 0.998 (0.506) |
| Reflections used in refinement (#) | 64955 (2783) |
| Reflections used for R-free (#) | 3165 (128) |
| R-work | 0.1969 |
| R-free | 0.2214 |
| Number of non-hydrogen atoms (#) | 1190 |
| macromolecules | 1018 |
| ligands | 16 |
| solvent | 156 |
| Protein residues (#) | 129 |
| RMS(bonds) | 0.027 |
| RMS(angles) | 2.2 |
| Ramachandran favored (%) | 98.43 |
| Ramachandran allowed (%) | 1.57 |
| Ramachandran outliers (%) | 0 |
| Rotamer outliers (%) | 0.93 |
| Clashscore | 5.44 |
| Average B-factor | 14.39 |
| macromolecules | 10.93 |
| ligands | 16.51 |
| solvent | 36.77 |

*Ab initio* phasing was performed by automatically placing idealized helical triple-alanine fragment followed by density modification (**Figure 1E, Materials and Methods**). A single helical fragment of just three alanine residues was sufficient to determine the entire lysozyme structure. After automatic placement in *PHASER*^29^, an interpretable map was produced following 144 rounds of dynamic density modification^18^, resulting in a map showing individually resolved atoms throughout the entire unit cell where all residues could be identified unambiguously even by eye (**Figure 2, Supplementary Movie 1 and 2**). The density-modified map (E, PHI) was similar to the final (2mF_o_-DF_c_, PHI) map after refinement. A complete model was automatically built into the density modified map given only the sequence^19^ and without consulting their known structures (**Figure 2**). For this structure, two C-terminal residues, and several solvent-exposed sidechains were either partially or absent in the map, even after the final refinement. They were also poorly resolved in X-ray investigations of triclinic lysozyme at similar resolutions^30^. The final models were refined^31^ using electron scattering factors.

**Figure 2.**
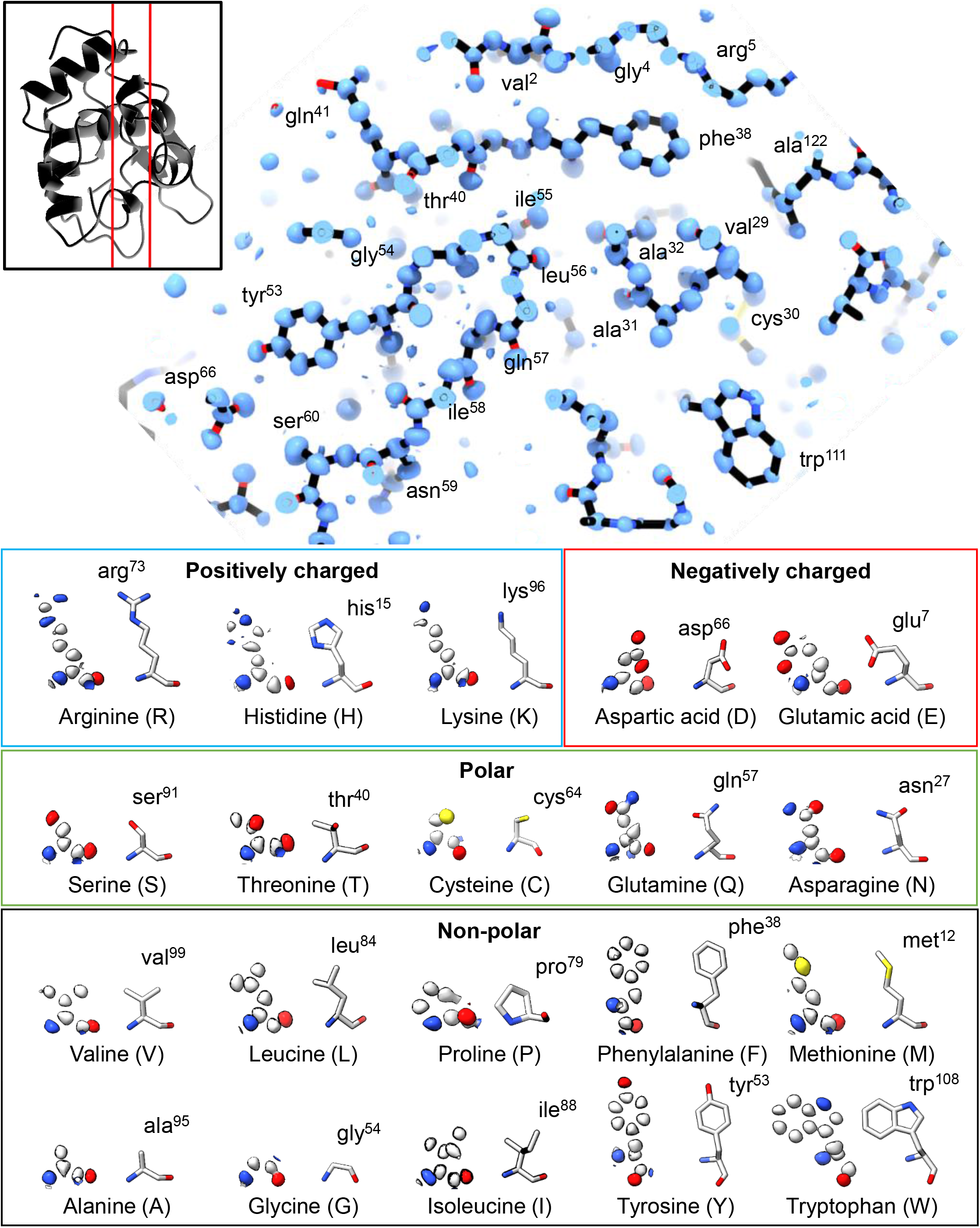
*Ab initio* structure of triclinic lysozyme at 0.87 Å resolution. **(Top)** A slice through the final structure of triclinic lysozyme as black sticks with the density modified map using normalized structure factors shown in blue. The location of the slice through the final structure is indicated by the inset on the top left. **(Bottom)** Examples of 20 representative amino acids from the final structure, one of each kind, are displayed with their normalized structure factor map at 0.87 Å resolution from density modification.

The results from lysozyme using electron counted MicroED data were very promising. However, it was not entirely surprising that sub-Ångstrom data could be determined *ab initio* even with the larger size of lysozyme compared with small molecules or small peptides. To test if other electron counted MicroED data would perform similarly, this approach was tested again using a sample of the serine protease, proteinase K, which is much larger and more representative of the average size of a globular protein in the cell. The crystals were grown in batch and the sample preparation was very similar to the approach used for the triclinic lysozyme microcrystals. The crystals were identically milled and screened (**Supplementary Figure 2**). Four proteinase K crystals were found on a single grid that met the selection criteria. MicroED data was collected using an overall similar approach using electron counting (**Materials and Methods**). It was similarly found that essentially all pixel values fell within the linear range of the detector and are still well below the threshold for damaging the detector (**Supplementary Figure 1, Materials and Methods**).

The data were indexed, integrated, and scaled using the same software as used for the triclinic lysozyme data. The space group was determined to be *P* 4 2 2 with a unit cell of, (a = b, c) (Å) = (67.08, 106.78), (α = β = γ) (°) = (90). Two of the four proteinase K lamellae integrated with lower internal consistency and were therefore discarded. A high-resolution cutoff was applied using the same criteria at 1.5 Å, and the overall completeness of the final dataset was 98.8% (**Supplementary Tables 2 & 3, Supplementary Figure 4**). The overall statistics for these merged crystals were better than any prior investigations on the same microscope using a CMOS^22^.

*Ab initio* phasing of larger proteins below atomic resolution is more challenging. The same approach was used as for lysozyme for phasing, namely automatically placing idealized helices and then extending the solution using density modification. Four 14 residue-long ideal alanine fragments were automatically placed using *PHASER*^29^. This solution was extended using multiple rounds of chain tracing and density modification^32^, similarly to the procedure implemented by ARCIMBOLDO_LITE^32,33^. The correct space group was determined by initially placing the fragments in all possible space groups for this point group, and then attempting to trace the best solutions. The solution in space group *P* 4_3_ 2_1_ 2 successfully built nearly the entire protein backbone. In the final round, this procedure traced 253 of 278 alanine residues (**Supplementary Table 2, Supplementary Figure 5**), and the map showed clear side-chain densities. The model was completed automatically^19^ from the backbone trace given only the sequence. The structure was similarly refined^31^ using electron scattering factors.

*Ab initio* phasing was successful for both lysozyme and proteinase K following a similar approach and using electron counted data. The lysozyme structure was determined from an idealized alanine helical fragment of only 15 atoms, corresponding to less than 1.5% of the structure. The maps calculated from only this small fragment showed density only around the placed atoms and uninterpretable noise elsewhere (**Figure 1E**) —not even the correct sidechains for the three placed residues could be reasonably modeled at this stage. However, the entire protein structure along with solvent ions and water molecules were individually resolved after density modification (**Figure 2**). The proteinase K structure was determined using four idealized helices, each made of fourteen alanine residues, or 280 atoms (**Materials and Methods, Supplementary Figure 5**), corresponding to about 13% of the structure. This solution also required density modification after placing the initial fragments. Its lower resolution and larger size meant that this structure required multiple rounds of chain tracing and density modification but was also determined *ab initio* using electron counting data. This procedure was repeatedly attempted using prior non-counting datasets but without success. The improved data quality from the electron counted data was critical to *ab initio* phasing macromolecular crystals using MicroED data.

This study demonstrated that MicroED data can be collected using a Falcon 4 direct electron detector in counting mode. Previous attempts at collecting MicroED data using counting on a Falcon 3 direct electron detector failed because the Falcon 3 detector is slower (40Hz) and had a much smaller linear range of only about 1 electron per pixel per second^16^. Therefore, the Falcon 3 was previously used in integrating (or linear) mode where it operates similarly to other CMOS-based detectors. In the current study, the Falcon 4 direct electron detector was used, which is capable of recording images at 250Hz. Combining the faster frame rate with an ultra-low dose spread over a long exposure meant that single electron events could be counted and the recorded intensities accurately integrated.

The data presented here were collected from crystal lamellae that were milled to an ideal thickness to match the mean free path of the microscope that was used^31^. Milling crystals can either be used to make large crystals into smaller ones^11,21,34,35^ to remove material that embeds small crystals and prevents accessing them for MicroED investigation, such as membrane proteins grown in viscous bicelles^23^ or LCP^36^. Regardless of the reason, milling crystals into an ideal thickness is good practice and recommended for extracting the most accurate data^22^. The lysozyme crystals milled in the current study diffract to sub-Ångström resolutions even after being milled by a gallium ion beam. This suggests that milling crystals does not compromise the sample or result in lower resolution. Indeed, the viability of collecting sub-Ångström data from crystalline lamellae further solidifies ion-beam milling as the preferred approach for preparing macromolecular crystals for MicroED experiments, since the crystals and surrounding material can be shaped and thinned to ideal thicknesses that match the inelastic mean free path for any accelerating voltage^22^. Importantly, at a mere 300nm, these lamellae matched the mean free path of a 300kV electron minimizing the probability of multiple scattering events and likely contributing the greater accuracy in the data.

The results presented here open the door for future investigations of phasing macromolecular protein crystals at or beyond near-atomic resolutions using MicroED data. Attempting to apply direct methods as implemented in *e*.*g*. Shake N Bake^37,38^ or SHELX^39,40^ are exciting avenues for future investigations since not all proteins will have helices to place for the starting phases. Further developments in *ab initio* phasing of MicroED data could be used to both automate the process and extend the necessary resolution to cover more challenging structures, such as membrane proteins that do not routinely reach resolutions better than about 3 Å. Regardless, the ability to probe membrane protein structure by MicroED is advantageous as electrons can probe the charge properties in the sample^41^. The quality of MicroED data obtained by counting would likely be further improved with the addition of an energy filter since this addition made large differences in the quality previously obtained on integrating detectors^17,41–43^. Given the importance of phase improvement in traditional X-ray crystallography^44^, the successful application of density modification algorithms compatible with MicroED data will be of critical importance for solving structures without known homologs or very difficult structures at lower resolutions.

## Materials and Methods

### Materials

Hen egg-white lysozyme (*Gallus gallus*) and Proteinase K (*Engyodontium album*) were purchased from Sigma-Aldrich (St. Louis, MO) and used without further purification. Sodium acetate pH 4.5, MES-NaOH pH 6.5, calcium chloride, and sodium nitrate stock solutions were used directly from Hampton (Viejo, CA) crystallization kits and diluted using milli-Q water as needed. The magic triangle kit was purchased from Hampton (Viejo, CA).

### Crystallization of triclinic lysozyme

Crystals were prepared similarly to the protocol originally detailed by Legrand et al.^45^, and then subsequently described by Heijna et al.^46^. Hen egg-white lysozyme was dissolved in 0.2 M NaNO_3_ 0.05 M sodium acetate pH 4.5 to a concentration of 10 mg / mL. A total volume of approximately 0.5 mL was prepared in a cold room and vortexed at the maximum setting for approximately 1 minute immediately after mixing. The tube containing this mixture was left at 4° C overnight. The next morning the tube was observed to be nearly filled with an opaque, white suspension. The sample was removed from the cold room, sealed with parafilm, and left on a benchtop at room temperature (approximately 23° C) for one week. After this time, very large, clear crystals accumulated on the bottom of the tube, and the rest of the liquid appeared transparent. Small (1 μL) aliquots from the center of the liquid that appeared clear by eye were found to contain a slurry of small, irregularly shaped crystals when viewed under a light microscope.

### Crystallization of proteinase K

Proteinase K was crystallized as described^47^. Protein powder was dissolved at a concentration of 40 mg / mL in 20 mM MES–NaOH pH 6.5. Crystals were formed by mixing a 1:1 ratio of protein solution and a precipitant solution composed of 0.5 M NaNO_3_, 0.1 M CaCl_2_, 0.1 M MES–NaOH pH 6.5 in a cold room at 4 °C. Microcrystals grew overnight.

### Grid preparation

Quantifoil Cu 200 R 2/2 holey carbon TEM grids were glow-discharged for 30 s at 15 mA on the negative setting immediately before use. Grids were loaded into a Leica GP2 vitrification robot. The robot sample chamber was loaded with filter paper and set to 4 °C and 95% humidity for 1 h before use. 3 μL of protein crystals from the center of either the proteinase K or lysozyme tubes were applied to the carbon side of the glow-discharged grid and allowed to incubate for 30 s. Grids were then gently blotted from the back for 20 s. For lysozyme, the grids were then immediately plunged into super-cooled liquid ethane. For proteinase K, 3 μL of 0.25M I3C 0.5 M NaNO_3_, 0.1 M CaCl_2_, 0.1 M MES–NaOH pH 6.5 was added as described^48^. The proteinase K grids were blotted once more from the back for 20 s and then immediately plunged into liquid ethane. Vitrified grids were stored at liquid nitrogen temperature before further experiments.

### Focused ion-beam and scanning electron microscopy

The vitrified grids were inserted into autogrid clips at liquid nitrogen temperature. After clipping, the grids were loaded into a cryo-transfer shuttle and inserted into a Thermo-Fisher Aquilos dual-beam FIB/SEM operating at liquid nitrogen temperatures. The samples were covered with first fine, then rough, coats of sputtered platinum immediately after loading. An additional layer of protective platinum was added using the gas injection system (GIS). GIS platinum coating was conducted in the mapping position at a working distance of 12 mm. In this way, an approximately 1 μm thick platinum layer was slowly deposited over 30 s and continually monitored using the electron beam.

Whole grid montages of each grid were taken at low magnification using the MAPS software (Thermo-Fisher). Crystals on the vitrified grids were identified in the FIB/SEM such that each crystal was not within 5 μm of a grid bar, not within three grid squares of the edge, and not within 25 μm of another selected crystal (**Figure 1A, Supplementary Figure 2A**). Twenty such crystals across two grids were prepared over two days for lysozyme and five proteinase K crystals on one grid were prepared in one day. Identified crystals were brought to the eucentric position and inspected in both the electron and ion beams. Milling was conducted as described^12^. Briefly, each crystal was roughly milled to approximately 3 μm thick and 5–10 μm wide using an ion-beam current of 300 pA and the standard rectangular milling patterns. Each crystal was then finely milled to a thickness of approximately 500 nm thick and 5–10 μm wide using an ion-beam current of 100 pA. Finally, each lamella was polished using an ion-beam current of 10 pA to a thickness of approximately 300 nm and width of 3–8 μm using a cleaning cross-section. This thickness was found to be optimum for experiments at this accelerating voltage^22^. Each of these steps is typically conducted in 1 – 5 min intervals using rectangular patterns with at most a 1 μm height dimension to reduce the influence of sample drift. The sample is imaged in the ion beam using a current of 1.5 pA between milling steps to realign as needed and to assess the quality and thickness of the finished lamella.

### Setting up the Falcon 4 for diffraction experiments

The dose protector that prevents the operation of this detector while in diffraction mode was disabled by a service engineer before experiments. The Falcon 4 direct electron detector internally operates at 250 frames per second, but every 32^nd^ frame window is used to reset the detector and does not result in useful data. Owing to bandwidth limitations, the camera furthermore accumulates at least seven raw frames before transmitting their summed image to the controlling computer, which means that the user can obtain no more than ∼35 images per second. For the data presented here, the frames were internally summed to correspond to 0.5 s wedges (119 raw frames; **Supplementary Figure 1**).

The damage threshold for this system is described by a deterioration of the detector quantum efficiency (DQE) due to radiation damage over time. Specifically, the DQE for each pixel will decrease by 10% after a total exposure of 1.5×10^9^ electrons. Ultimately, the smaller C2 aperture of 50 μm was chosen, with a spot size of 11, and a beam diameter of 20 or 25 μm which corresponded to a total exposure of ∼0.64 – 1.0 e^-^ Å^-2^. The typical counts on the detector were within the linear region and a typical pixel experienced a smaller exposure than typical single-particle movies (**Supplementary Figure 1**). None of our measurements fell below a DQE of 0.6.

### MicroED data collection

Grids containing milled protein crystals were rotated such that the TEM rotation axis was 90° from the FIB/SEM’s milling axis, and then loaded into a cryogenically cooled Thermo-Fisher Titan Krios 3Gi transmission electron microscope operating at an accelerating voltage of 300 kV. Low magnification montages of each grid were collected at a magnification of 64 × and used to locate the milled lamellae on each grid. Each lamella was brought to its eucentric position before data collection. MicroED data were collected by continuously rotating the stage at a rate of approximately 0.15° / s or 0.2° / s for 420 s, covering a total rotation range of approximately 63° or 84°, respectively. This typically spanned the real space wedge corresponding to -31.5° to +31.5°, or -42° to +42°. Data were collected using a 50 μm C2 aperture, a spot size of 11, and a beam diameter of 20 or 25 μm. Under these conditions, the total exposure per dataset was approximately 1.0 e^-^ Å^-2^ or 0.64 e^-^ Å^-2,^ respectively. Diffraction data were isolated from a small area from the middle of each lamella of approximately 2 μm in diameter at the specimen level using the 100 μm selected area aperture to remove unwanted background noise. All data were collected using twofold binning and internally summed such that each image records a 0.5 s exposure spanning approximately 0.075– 0.1° of rotation. In this way, each image stack contained 840 images, the last of which was discarded. A single sweep of continuous rotation MicroED data was collected from each lysozyme lamella. For proteinase K, two sweeps were collected—a high-resolution dataset at a nominal camera distance of 960 mm, and then a subsequent low-resolution dataset collected identically, but at the longest possible nominal camera distance of 4300 mm. The post-column magnification on this system is 1.81×. The low-resolution pass was conducted after the high-resolution pass covering a resolution range from approximately 60–6 Å.

### MicroED data processing

Movies in MRC format were converted to SMV format using a parallelized version of the MicroED tools^12^ (https://cryoem.ucla.edu/downloads). Each dataset was indexed and integrated using XDS. All datasets were scaled using XSCALE^27^ and xscale_isocluster^49^. For both crystals, the space group was verified using POINTLESS ^50^. For lysozyme, the data were merged without scaling using AIMLESS ^50^, the subsequent intensities were converted to amplitudes in CTRUNCATE ^26^, normalized structure factors were calculated using ECALC ^26^, and a 5% fraction of the reflections assigned to a free set using FREERFLAG software packages distributed in the CCP4 program suite^26^.

### Phasing

The lysozyme data could be phased using either FRAGON^51^, or PHASER^29^ followed by ACORN^18,52^. An initial phasing solution was achieved using example parameters from the ACORN documentation (http://legacy.ccp4.ac.uk/html/acorn.html#example9). Here, a small fragment of idealized alpha helix is used for molecular replacement, and then the best solution is subjected to density modification. In this way, the number of atoms was systematically lowered in the idealized helix from the initial 50 and were able to achieve a phasing solution with as few as 15 total atoms. Using fragments smaller than 20 atoms required placement using PHASER rather than the internal ACORN procedure. The resolution limits were also tested using the same 50 atom fragment, and the structure was solved using the same procedure up to 1.15 Å resolution. The lysozyme model could also be solved in FRAGON starting from a penta-alanine fragment, the smallest fragment allowed in the CCP4I2 interface^53^.

Additional tests were set up to explore the limits of our solution under different circumstances. First, a 10 alanine (50 atoms) helix was used for molecular replacement at resolutions worse than 0.87 Å. A single helix could be placed with high accuracy to at least 1.5 Å resolution. Using this solution as the initial phases for the ACORN density modification procedure using the data to 1.0 Å resulted in a map nearly identical to the one that had the helix placed using the entire resolution range. However, individual atoms were no longer resolved after density modification with data worse than 1.15 Å resolution. No resolution extension features were used in any runs of ACORN.

For proteinase K, ideal helices were placed using PHASER^29^. Four copies of the idealized 14 amino acid alanine helix were placed in space groups #89 - #96. SHELXE was run by using the merged intensities using the MTZ2HKL utility. Surprisingly, using the unmerged intensities directly from XSCALE did not result in a successful trace. Of the attempts of SHELXE from all the space groups, only #96 (*P* 4_3_ 2_1_ 2) resulted in a well-traced structure with clear sidechain density. From here, 1, 2, and 3 copies of 14 long alpha-helices were placed in space group #96. This was attempted again using 10 amino acid long helices, where only the search for 4 copies resulted in a convincing solution. SHELXE was run with or without the “-q” flag to first search for helix shapes during the chain tracing. The first solution from 4 helices in space group #96 used the SHELXE command line: “shelxe 1.pda –s0.4 –a30.” Here, the default 10 rounds of density modification are followed by standard chain tracing. This is repeated 30 times. Among these trials, only the four placed 14-long helices gave an obvious solution after chain tracing. However, the solution in space group #96 with 3 helices placed was also able to give a similar solution upon adding the –o option to trim away the low CC amino acids, -q to search for helical shapes, and increasing the chain tracing rounds to 50 or 100.

For both lysozyme and proteinase K after the last round of density modification, the protein was built automatically by BUCCANEER^19^. For lysozyme, the entire protein was built into the map produced by ACORN^18^ except for two terminal residues that were not resolved upon inspection of the map in Coot^54^. For proteinase K, the traced backbone from SHELXE was used as a starting fragment for BUCCANEER. In both cases, electron scattering factors were used for the maps. The built structures were refined in REFMAC^31^ using electron scattering factors calculated by the Mott-Bethe formula. Initial refinements used isotropic atomic displacement (B) factors for individual atoms and waters were added automatically. Refinement was always followed by manual curation of the model using Coot. For lysozyme, NO_3_ ions were found in multiple locations that were not adequately modeled by single water molecules. For proteinase K, two I3C molecules with low occupancies were identified after the structure was entirely built and placed manually in Coot using the fragment code, I3C. Hydrogen atoms were added to the lysozyme model in their riding positions for the final rounds of refinement. Once the model was completely built, the models were refined again using anisotropic atomic displacement parameters for all but the hydrogen atoms.

## Figure and table legends

**Supplementary Table 1.** Merging statistics for triclinic lysozyme.

| Resolution | # Observed | # Unique | # Possible | Multiplicity | Completeness | R obs | R expected | I/Sigma | R meas | CC 1/2 | R-p.i.m. |
| --- | --- | --- | --- | --- | --- | --- | --- | --- | --- | --- | --- |
| 1.87 | 76398 | 7399 | 7501 | 10.3 | 98.60% | 20.30% | 19.50% | 14.56 | 21.30% | 98.30 | 6.2% |
| 1.49 | 79348 | 7268 | 7300 | 10.9 | 99.60% | 29.30% | 31.50% | 11.28 | 30.80% | 96.30 | 9.1% |
| 1.3 | 87189 | 7490 | 7500 | 11.6 | 99.90% | 41.00% | 63.40% | 8.56 | 43.00% | 92.20 | 12.4% |
| 1.18 | 87995 | 7463 | 7472 | 11.7 | 99.90% | 57.20% | 111.30% | 6.67 | 60.00% | 83.70 | 17.3% |
| 1.1 | 65854 | 6840 | 6980 | 9.6 | 98.00% | 55.20% | 173.80% | 5.43 | 58.40% | 81.10 | 18.5% |
| 1.03 | 62848 | 7720 | 8027 | 8.1 | 96.20% | 60.00% | 247.40% | 3.98 | 64.10% | 77.50 | 22.1% |
| 0.98 | 44775 | 6521 | 7202 | 6.8 | 90.50% | 73.80% | 87.60% | 2.48 | 79.80% | 67.90 | 29.0% |
| 0.94 | 33255 | 5631 | 6917 | 5.9 | 81.40% | 97.90% | 128.20% | 1.65 | 106.80% | 50.40 | 41.6% |
| 0.9 | 26371 | 6012 | 8240 | 4.3 | 73.00% | 117.50% | 238.00% | 1.02 | 132.80% | 31.30 | 59.9% |
| 0.87 | 5378 | 2644 | 7147 | 2.0 | 37.00% | 102.10% | 136.60% | 0.65 | 140.60% | 14.60 | 94.6% |
| total | 569411 | 64988 | 74286 | 8.7 | 87.50% | 23.60% | 27.10% | 6.23 | 24.80% | 99.00 | 7.3% |

**Supplementary Table 2.** MicroED crystallographic table of of proteinase K.

| MicroED Data and Structure |  |
| --- | --- |
| Proteinase K |  |
| Wavelength (Å) | 0.0197 |
| # of crystals (#) | 2 |
| Resolution range (Å) | 43.35 - 1.5 (1.554 - 1.5) |
| Space group | P 4<sub>3</sub> 2<sub>1</sub> 2 |
| Unit cell (a, b, c) (Å) | 67.08, 67.08, 106.78 |
| (α, β, γ) (°) | 90, 90, 90 |
| Total reflections (#) | 416133 (36794) |
| Unique reflections (#) | 39347 (3683) |
| Multiplicity | 10.6 (9.9) |
| Completeness (%) | 98.87 (94.41) |
| < I/σ(I) > | 5.65 (1.12) |
| Wilson B-factor | 13.16 |
| R-merge | 0.277 (1.508) |
| R-meas | 0.291 (1.59) |
| R-pim | 0.08711 (0.4921) |
| CC<sub>1/2</sub> | 0.989 (0.31) |
| CC* | 0.997 (0.688) |
| Reflections used in refinement (#) | 39301 (3683) |
| Reflections used for R-free (#) | 2001 (210) |
| R-work | 0.1495 |
| R-free | 0.2046 |
| Number of non-hydrogen atoms (#) | 2300 |
| macromolecules | 2029 |
| ligands | 37 |
| solvent | 234 |
| Protein residues (#) | 279 |
| RMS(bonds) | 0.015 |
| RMS(angles) | 1.89 |
| Ramachandran favored (%) | 97.11 |
| Ramachandran allowed (%) | 2.89 |
| Ramachandran outliers (%) | 0 |
| Rotamer outliers (%) | 0.94 |
| Clashscore | 4.99 |
| Average B-factor | 16.12 |
| macromolecules | 14.19 |
| ligands | 46.67 |
| solvent | 28.02 |

**Supplementary Table 3.** Merging statistics for proteinase K.

| Resolution | # Observed | # Unique | # Possible | Multiplicity | Completeness | R-obs | R- expected | I/Sigma | R meas | CC 1/2 | R-p.i.m. |
| --- | --- | --- | --- | --- | --- | --- | --- | --- | --- | --- | --- |
| 3.23 | 47759 | 4205 | 4257 | 11.35767 | 98.80% | 15.60% | 18.30% | 12.2 | 16.30% | 98.6 | 6.4% |
| 2.57 | 40343 | 3972 | 3999 | 10.15685 | 99.30% | 19.90% | 19.90% | 9.73 | 21.00% | 98.2 | 8.9% |
| 2.24 | 41742 | 4007 | 4036 | 10.41727 | 99.30% | 27.90% | 25.90% | 7.67 | 29.40% | 96.6 | 10.4% |
| 2.04 | 40455 | 3850 | 3873 | 10.50779 | 99.40% | 32.80% | 31.50% | 6.59 | 34.50% | 95.2 | 13.4% |
| 1.89 | 42628 | 4027 | 4047 | 10.58555 | 99.50% | 42.20% | 42.00% | 5.18 | 44.40% | 92.0 | 18.2% |
| 1.78 | 40986 | 3848 | 3869 | 10.65125 | 99.50% | 57.80% | 66.10% | 3.8 | 60.70% | 84.3 | 24.7% |
| 1.69 | 42082 | 3935 | 3955 | 10.69428 | 99.50% | 78.00% | 111.20% | 2.73 | 81.90% | 74.5 | 31.7% |
| 1.55 | 86252 | 8052 | 8089 | 10.71187 | 99.50% | 110.80% | 192.50% | 1.82 | 116.40% | 55.4 | 38.7% |
| 1.5 | 33884 | 3451 | 3637 | 9.818603 | 94.90% | 152.50% | 491.40% | 1.14 | 160.80% | 29.7 | 49.2% |
| total | 416131 | 39347 | 39762 | 10.57593 | 99.00% | 27.80% | 35.40% | 5.36 | 29.20% | 98.8 | 8.7% |

**Supplementary Figure 1.**
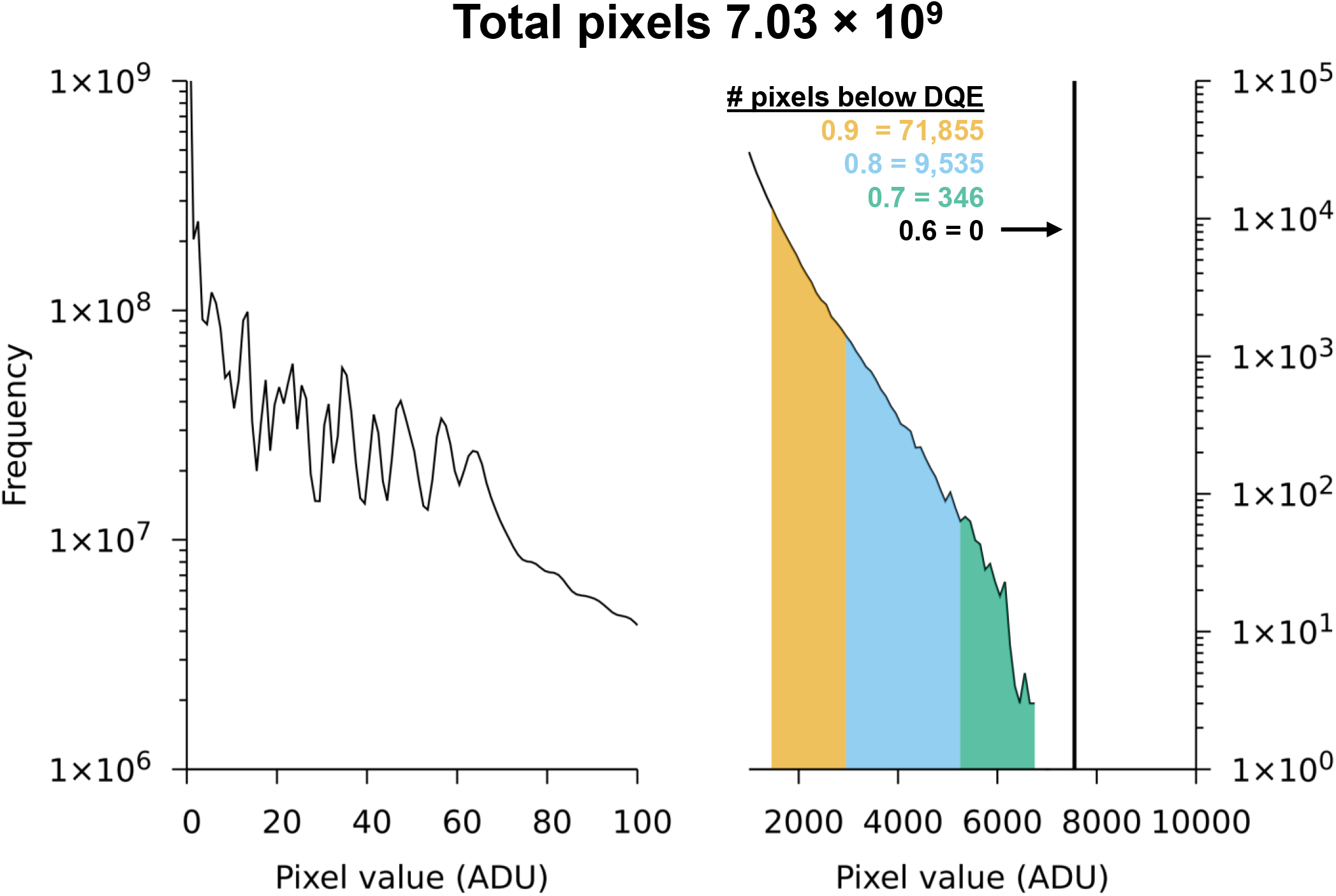
Logarithmic histograms of intensities from both proteinase K datasets. (Left) low-count region showing resolved individual counts. (Right) Higher count region showing the distribution of counts below the indicated theoretical DQE threshold. No pixel counts fell below a DQE value of 0.6 for these higher dose datasets.

**Supplementary Figure 2.**
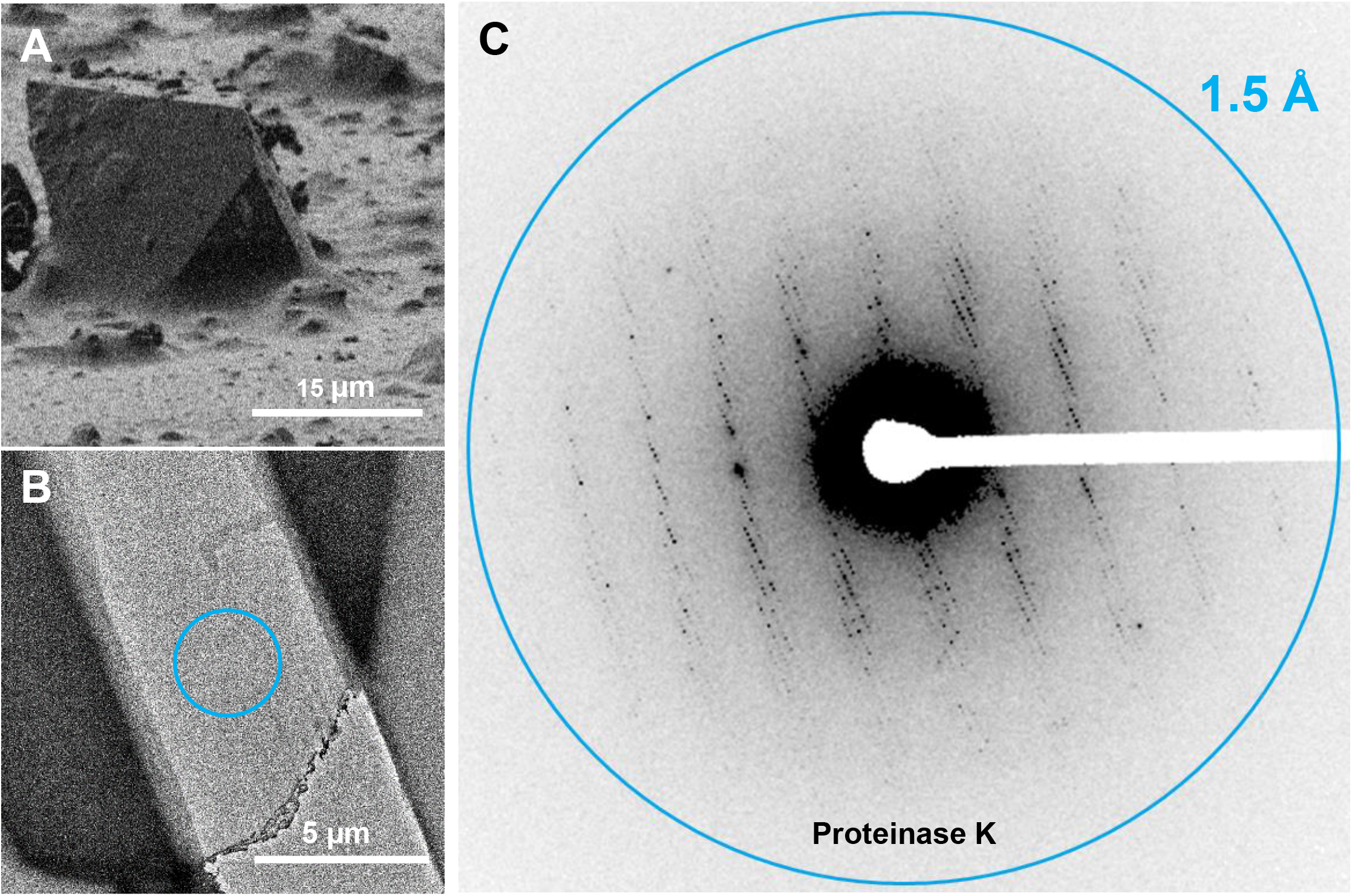
Electron counted MicroED data from milled lamellae of proteinase K. **(A)** A typical proteinase K microcrystal imaged using the focused ion beam. **(B)** A thin, milled lamella from (A) identified in the TEM. **(C)** MicroED data collected in counting mode on a Falcon 4 direct electron detector from a proteinase lamella.

**Supplementary Figure 3.**
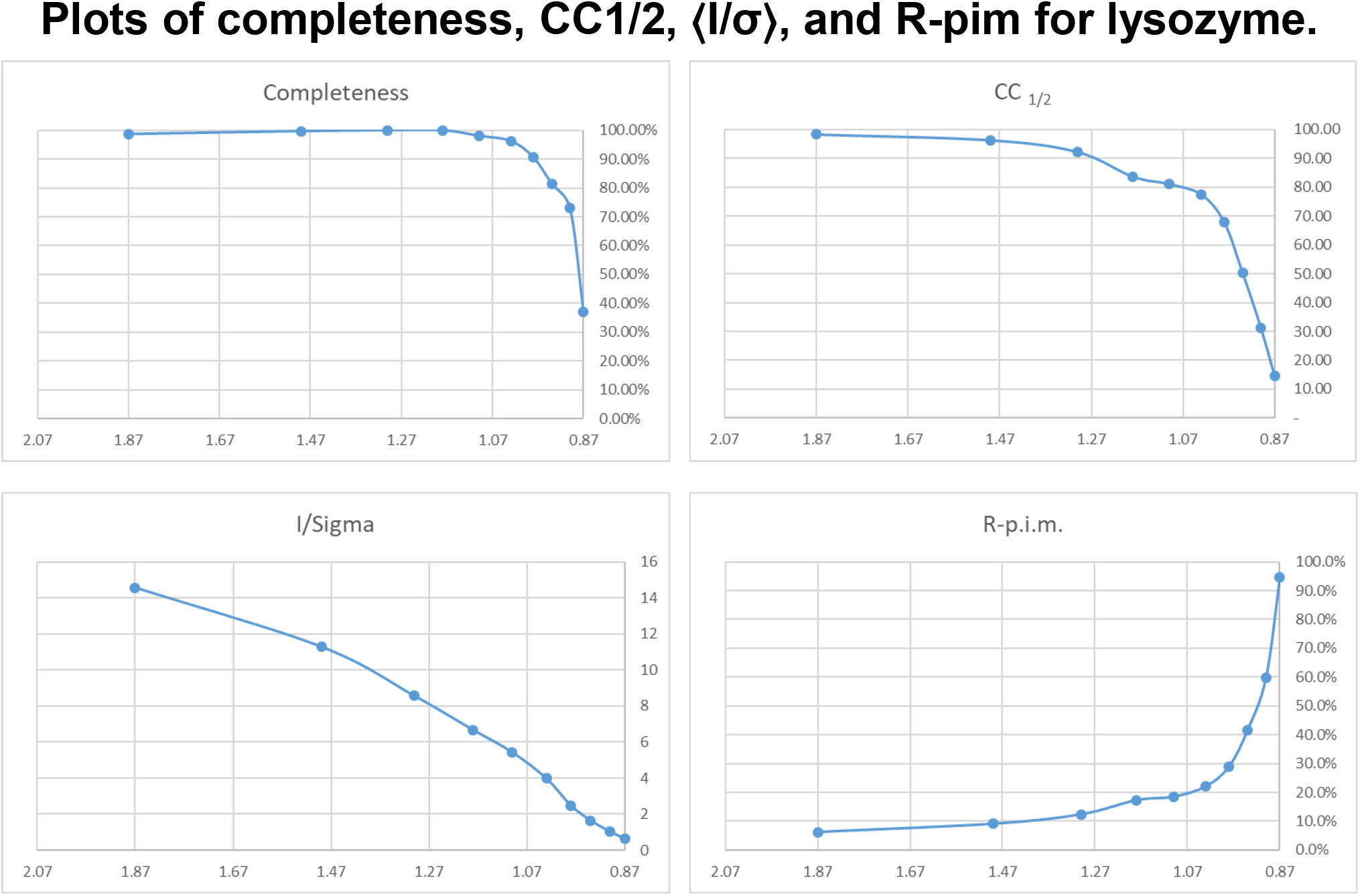
Plots of completeness, CC_1/2_, ⟨I/σ⟩, and R-pim for lysozyme.

**Supplementary Figure 4.**
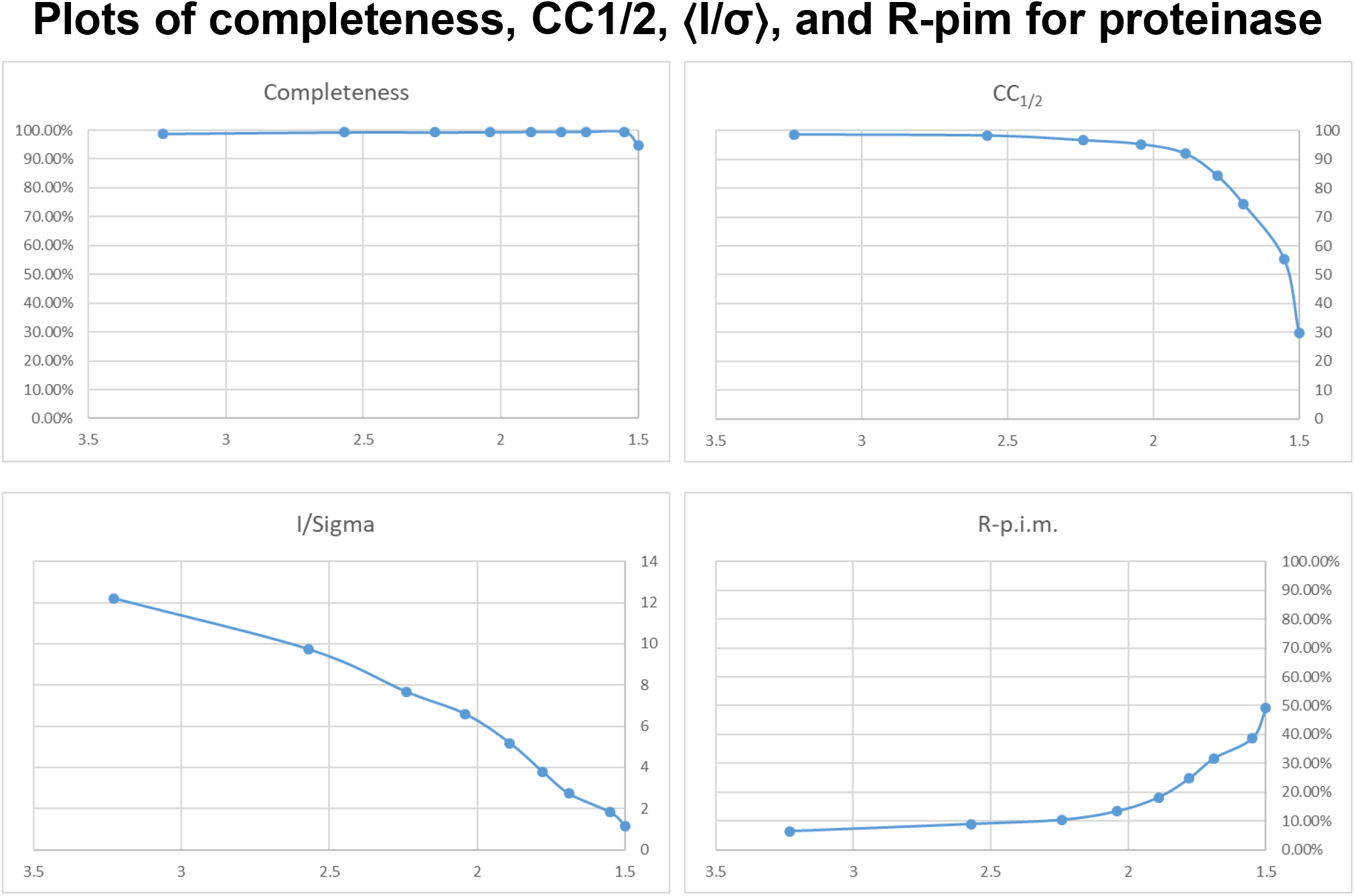
Plots of completeness, CC_1/2_, ⟨I/σ⟩, and R-pim for proteinase K.

**Supplementary Figure 5.**
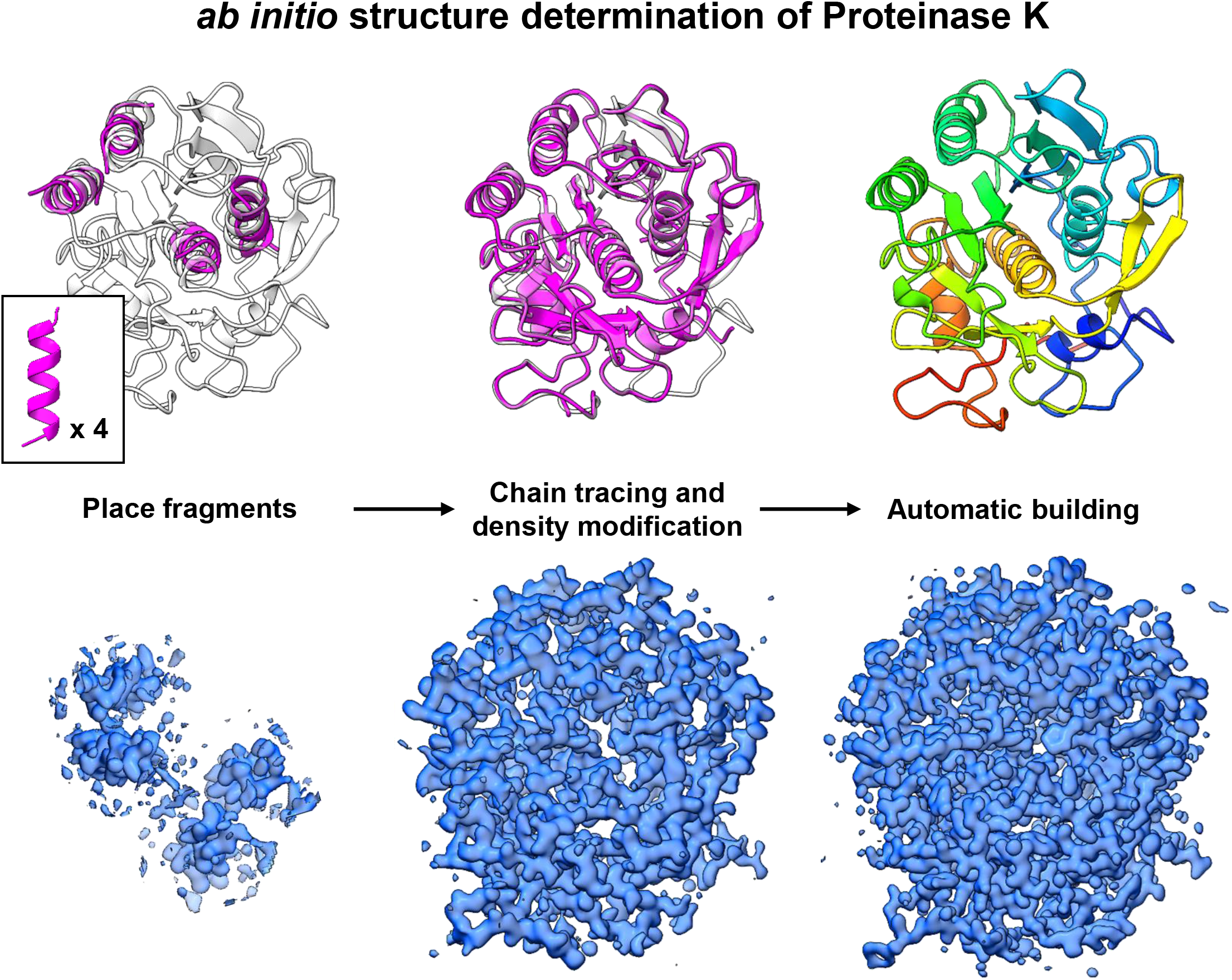
*Ab initio* structure of Proteinase K at 1.5 Å resolution. **(Top)** The progression of the models during the phasing process. Four, 14 alanine fragments with idealized geometry were chosen as the starting fragments. The fragments were placed using molecular replacement. From this initial placement, repeated density modification and chain tracing were conducted. The chain traced model was used as an input for automatic building, and the entire protein structure was completed automatically. **(Bottom)** maps at the intermediate stages of the phasing process. 2mF_o_-DF_c_ maps are in white and contoured at the 1.0 σ level, whereas the mF_o_-DF_c_ maps are contoured at the +/-3.0 σ levels and presented in green and red, respectively.

**Supplementary Movie 1**. A rotation movie showing the normalized structure factor maps after density modification over the final structure of lysozyme at 0.87 Å resolution. The movie is sliced to a width of 8 Å.

**Supplementary Movie 2**. A rotation movie showing the normalized structure factor maps after density modification over the final structure showing a region rich with hydrophobic packing interactions. Tryptophan, isoleucine, tryptophan sandwich are shown with several other tryptophan residues flanking this region. The movie is sliced to a width of 8 Å.

## Figure preparation

Figures were prepared using ChimeraX and Pymol, then arranged in PowerPoint. Tables were arranged in Excel. Images were cropped around areas of interest, and the brightness and contrast were adjusted in FIJI^55^.

## Software availability

Software tools that convert summed or unsummed MRC stacks to Super Marty View or TIFF are available at https://cryoem.ucla.edu/MicroED.

## Funding information

This study was supported by the National Institutes of Health P41GM136508. The Gonen laboratory is supported by funds from the Howard Hughes Medical Institute.

## Acknowledgments

The authors would like to thank Drs. Lingbo Yu and Bart Buijsse (Thermo-Fisher) for information and discussions about the Falcon 4 and counting mode. Coordinates for triclinic lysozyme and proteinase K structures have been deposited to the PDB, and maps have been deposited to the EMDB.

## Notes

### Competing Interest Statement

The authors have declared no competing interest.

## References

1. Nannenga, B. L. & Gonen, T. The cryo-EM method microcrystal electron diffraction (MicroED). Nature methods 16, 369–379 (2019).

2. Nannenga, B. L., Shi, D., Leslie, A. G. & Gonen, T. High-resolution structure determination by continuous-rotation data collection in MicroED. Nature methods 11, 927–930 (2014).

3. Arndt, U. W. & Wonacott, A. J. Rotation method in crystallography. (North-Holland Pub. Co., 1977).

4. Sawaya, M. R. et al. Ab initio structure determination from prion nanocrystals at atomic resolution by MicroED. Proc Natl Acad Sci USA 113, 11232–11236 (2016).

5. Hiroaki, Y. et al. Implications of the Aquaporin-4 Structure on Array Formation and Cell Adhesion. Journal of Molecular Biology 355, 628–639 (2006).

6. Maeda, S. et al. Structure of the connexin 26 gap junction channel at 3.5 Å resolution. Nature 458, 597–602 (2009).

7. Wisedchaisri, G. & Gonen, T. Fragment-based phase extension for three-dimensional structure determination of membrane proteins by electron crystallography. Structure 19, 976–987 (2011).

8. Clabbers, M. T. B. et al. MyD88 TIR domain higher-order assembly interactions revealed by microcrystal electron diffraction and serial femtosecond crystallography. Nat Commun 12, 2578 (2021).

9. Xu, H. et al. Solving a new R2lox protein structure by microcrystal electron diffraction. Sci. Adv. 5, eaax4621 (2019).

10. Richards, L. S. et al. Fragment-based determination of a proteinase K structure from MicroED data using ARCIMBOLDO_SHREDDER. Acta Crystallogr D Struct Biol 76, 703–712 (2020).

11. Zhou, H., Luo, Z. & Li, X. Using focus ion beam to prepare crystal lamella for electron diffraction. Journal of Structural Biology 205, 59–64 (2019).

12. Martynowycz, M. W., Zhao, W., Hattne, J., Jensen, G. J. & Gonen, T. Collection of Continuous Rotation MicroED Data from Ion Beam-Milled Crystals of Any Size. Structure 27, 545-548.e2 (2019).

13. Clabbers, M. T. et al. Protein structure determination by electron diffraction using a single three-dimensional nanocrystal. Acta Crystallographica Section D: Structural Biology 73, 738–748 (2017).

14. Henderson, R. Overview and future of single particle electron cryomicroscopy. Archives of biochemistry and biophysics 581, 19–24 (2015).

15. Kuhlbrandt, W. The Resolution Revolution. Science 343, 1443–1444 (2014).

16. Hattne, J., Martynowycz, M. W., Penczek, P. A. & Gonen, T. MicroED with the Falcon III direct electron detector. IUCrJ 6, (2019).

17. Takaba, K., Maki-Yonekura, S., Inoue, S., Hasegawa, T. & Yonekura, K. Protein and Organic-Molecular Crystallography With 300kV Electrons on a Direct Electron Detector. Front. Mol. Biosci. 7, (2021).

18. Dodson, E. J. & Woolfson, M. M. ACORN2: New developments of the ACORN concept. Acta Crystallographica Section D: Biological Crystallography 65, 881–891 (2009).

19. Cowtan, K. The Buccaneer software for automated model building. 1. Tracing protein chains. Acta crystallographica section D: biological crystallography 62, 1002–1011 (2006).

20. Hattne, J., Shi, D., de la Cruz, M.J., Reyes, F. E. & Gonen, T. Modeling truncated pixel values of faint reflections in MicroED images. J Appl Crystallogr 49, 1029–1034 (2016).

21. Beale, E. V. et al. A Workflow for Protein Structure Determination From Thin Crystal Lamella by Micro-Electron Diffraction. Front. Mol. Biosci. 7, 179 (2020).

22. Martynowycz, M. W., Clabbers, M. T. B., Unge, J., Hattne, J. & Gonen, T. Benchmarking ideal sample thickness in cryo-EM using MicroED. 2021.07.02.450941 https://www.biorxiv.org/content/10.1101/2021.07.02.450941v1 (2021) doi:10.1101/2021.07.02.450941.

23. Martynowycz, M. W., Khan, F., Hattne, J., Abramson, J. & Gonen, T. MicroED structure of lipid-embedded mammalian mitochondrial voltage-dependent anion channel. Proc Natl Acad Sci USA 117, 32380–32385 (2020).

24. Hattne, J. et al. Analysis of Global and Site-Specific Radiation Damage in Cryo-EM. Structure 26, 759-766.e4 (2018).

25. Kabsch, W. XDS. Acta Crystallogr D Biol Crystallogr 66, 125–132 (2010).

26. Winn, M. D. et al. Overview of the CCP4 suite and current developments. Acta Crystallographica Section D: Biological Crystallography 67, 235–242 (2011).

27. Kabsch, W. Integration, scaling, space-group assignment and post-refinement. Acta Crystallogr D Biol Crystallogr 66, 133–144 (2010).

28. Karplus, P. A. & Diederichs, K. Linking Crystallographic Model and Data Quality. Science 336, 1030–1033 (2012).

29. McCoy, A. J. et al. Phaser crystallographic software. J Appl Crystallogr 40, 658–674 (2007).

30. Walsh, M. A. et al. Refinement of triclinic hen egg-white lysozyme at atomic resolution. Acta Crystallographica Section D: Biological Crystallography 54, 522–546 (1998).

31. Kovalevskiy, O., Nicholls, R. A., Long, F., Carlon, A. & Murshudov, G. N. Overview of refinement procedures within REFMAC5: utilizing data from different sources. Acta Crystallographica Section D: Structural Biology 74, 215–227 (2018).

32. Thorn, A. & Sheldrick, G. M. Extending molecular-replacement solutions with SHELXE. Acta Cryst D 69, 2251–2256 (2013).

33. Sammito, M. et al. Exploiting tertiary structure through local folds for crystallographic phasing. Nature Methods 10, 1099–1101 (2013).

34. Duyvesteyn, H. M. E. et al. Machining protein microcrystals for structure determination by electron diffraction. Proc Natl Acad Sci USA 115, 9569–9573 (2018).

35. Li, X., Zhang, S., Zhang, J. & Sun, F. In situ protein micro-crystal fabrication by cryo-FIB for electron diffraction. Biophys Rep 4, 339–347 (2018).

36. Martynowycz, M. W. et al. MicroED structure of the human adenosine receptor determined from a single nanocrystal in LCP. Proc Natl Acad Sci USA 118, e2106041118 (2021).

37. Miller, R., Gallo, S. M., Khalak, H. G. & Weeks, C. M. SnB: crystal structure determination via shake-and-bake. Journal of Applied Crystallography 27, 613–621 (1994).

38. Weeks, C. M. & Miller, R. Optimizing Shake-and-Bake for proteins. Acta Crystallographica Section D: Biological Crystallography 55, 492–500 (1999).

39. Sheldrick, G. M. A short history of SHELX. Acta Crystallographica Section A: Foundations of Crystallography 64, 112–122 (2008).

40. Usón, I. & Sheldrick, G. M. An introduction to experimental phasing of macromolecules illustrated by SHELX; new autotracing features. Acta Crystallographica Section D: Structural Biology 74, 106–116 (2018).

41. Yonekura, K. & Maki-Yonekura, S. Refinement of cryo-EM structures using scattering factors of charged atoms. Journal of Applied Crystallography 49, 1517–1523 (2016).

42. Gonen, T. et al. Lipid–protein interactions in double-layered two-dimensional AQP0 crystals. Nature 438, 633–638 (2005).

43. Yonekura, K., Maki-Yonekura, S. & Namba, K. Quantitative Comparison of Zero-Loss and Conventional Electron Diffraction from Two-Dimensional and Thin Three-Dimensional Protein Crystals. Biophysical Journal 82, 2784–2797 (2002).

44. Cowtan, K. Recent developments in classical density modification. Acta Crystallographica Section D: Biological Crystallography 66, 470–478 (2010).

45. Legrand, L., Riès-Kautt, M. & Robert, M. C. Two polymorphs of lysozyme nitrate: temperature dependence of their solubility. Acta Crystallogr D Biol Crystallogr 58, 1564–1567 (2002).

46. Heijna, M. C. R., van den Dungen, P. B. P., van Enckevort, W. J. P. & Vlieg, E. An Atomic Force Microscopy Study of the (001) Surface of Triclinic Hen Egg-White Lysozyme Crystals. Crystal Growth & Design 6, 1206–1213 (2006).

47. Masuda, T. et al. Atomic resolution structure of serine protease proteinase K at ambient temperature. Sci Rep 7, 45604 (2017).

48. Martynowycz, M. W. & Gonen, T. Ligand Incorporation into Protein Microcrystals for MicroED by On-Grid Soaking. Structure (2020).

49. Diederichs, K. Dissecting random and systematic differences between noisy composite data sets. Acta Cryst D 73, 286–293 (2017).

50. Evans, P. R. & Murshudov, G. N. How good are my data and what is the resolution? Acta Crystallogr D Biol Crystallogr 69, 1204–1214 (2013).

51. Jenkins, H. T. Fragon: rapid high-resolution structure determination from ideal protein fragments. Acta Crystallographica Section D: Structural Biology 74, 205–214 (2018).

52. Foadi, J. General concepts underlying ACORN, a computer program for the solution of protein structures. Crystallography Reviews 9, 43–65 (2003).

53. Potterton, L. et al. CCP4i2: the new graphical user interface to the CCP4 program suite. Acta Crystallographica Section D: Structural Biology 74, 68–84 (2018).

54. Emsley, P., Lohkamp, B., Scott, W. G. & Cowtan, K. Features and development of Coot. Acta Crystallographica Section D: Biological Crystallography 66, 486–501 (2010).

55. Schindelin, J. et al. Fiji: an open-source platform for biological-image analysis. Nat Methods 9, 676–682 (2012).

